# Systematic identification of oscillatory gene expression in single cell types

**DOI:** 10.1101/2025.09.02.673125

**Authors:** Alexis Weinreb, Manasa Basavaraju, Marc Hammarlund, Maxwell G. Heiman

**Author notes:** Correspondence (M. H.); (M. G. H.).

## Abstract

Many biological cycles are driven by oscillatory gene expression coordinated across cell types. For example, larval development in *Caenorhabditis elegans* involves coordinated cyclic changes in cell division, behavior, and growth, the latter requiring production of a structured extracellular matrix called the cuticle. Here, we combine single-cell RNA sequencing and novel computational approaches to identify oscillatory gene expression in individual cell types. We find that many cell types exhibit looping structures in PCA and UMAP space that correspond to transcriptional oscillations at each larval stage. Oscillatory gene expression is found in all cuticle-producing cell types, including glia, but not detected in neurons or muscle. We develop rigorous statistical approaches for *de novo* identification of oscillatory genes and cell types, yielding >5,000 genes. While many oscillatory genes relate to cuticle production, each cell type expresses largely distinct genes, suggesting that cuticle production is a patchwork of cell-type-specific programs. Finally, we derive a potential set of regulatory transcription factors that can explain coordinated oscillatory gene expression and find that shared upstream factors likely control gene timing across cell types. Together, our results suggest that shared regulators control cell-type-specific oscillatory gene expression, including in previously overlooked cell types such as glia.

## INTRODUCTION

Organisms undergo physiological cycles ranging from hours to weeks or months (e.g., circadian rhythms, estrous cycle, seasonal hibernation or migration). During such cycles, organismal-level changes arise from coordinated oscillatory gene programs in individual tissues. A simple model is that a time-keeping mechanism, called an oscillator, is shared across tissues and combines with cell-type-specific transcription factors to produce different transcriptional outputs in each tissue. Evidence from *Drosophila* circadian cycles supports this model (Meireles-Filho et al. 2014).

The larval cycle of *Caenorhabditis elegans* offers a powerful experimental system in which to dissect cell-type-specific features of oscillatory gene programs. During development, *C. elegans* proceeds through four stereotyped larval stages (L1, L2, L3, and L4) punctuated by molts. During each larval stage, the animal eats, grows, and executes behavioral programs to explore its environment. At molts, the animal enters lethargus, characterized by a quiescent state (sleep) alternating with bouts of activity. It also synthesizes a new cuticle – an apical extracellular matrix (aECM) that covers the entire body. After lethargus ends, the animal completes the molt by shedding its old cuticle like a snake skin. The timing of many cell biological events is coupled to progression through the larval cycle, for example hypodermal seam cells undergo cell division at a defined time in each larval stage. The larval cycle thus involves diverse, coordinated changes across cell types, including alterations in neuronal activity, cuticle synthesis, and cell cycle control. This is accompanied by widespread transient gene expression that repeats at each larval stage (Kim et al. 2013; Hendriks et al. 2014; Meeuse et al. 2020). These changes are controlled in part by the ancestral oscillator *lin-42*/Period, a conserved regulator of circadian rhythms, suggesting that insights from the *C. elegans* larval cycle may be generalizable across systems (Lazetic and Fay 2017; Spangler et al. 2024).

Synthesis of a new cuticle at the molting phase of each larval cycle is critical, as the cuticle is a complex structure that must protect the animal from its environment while allowing for growth, movement, and sensory perception. While most studies of cuticle synthesis have focused primarily on the skin (i.e., hyp7 and seam cells), we recently noted that some glial cells – specifically, a subtype called IL socket (ILso) glia – exhibit transient expression of an aECM protein involved in patterning a specialized cuticle covering for sensory neurons (Fung et al. 2023). These observations led to the hypothesis that some glia exhibit their own oscillatory gene program, distinct from that of the skin, and raised the question whether other unexplored cell types might do so as well.

Here, we use single-cell RNA sequencing to identify cell-type-specific oscillatory gene programs.

We find that socket glia exhibit oscillatory expression of hundreds of genes. Significant oscillatory patterns are found in other cuticle-producing cells, such as skin and pharynx, but are not evident in cells such as neurons and muscle, suggesting that cuticle synthesis is a major determinant of oscillatory gene expression. We develop a computational method for *de novo* identification of genes with pulsatile expression from single-cell RNA sequencing data, and use it to discover novel oscillatory genes expressed in glia, skin, pharynx, and other cell types. The largest families of pulsatile genes encode putative aECM proteins, consistent with a role in cuticle synthesis.

Surprisingly, most pulsatile genes are specific to small sets of cell types, suggesting previously unappreciated local differences in cuticle patterning. To discover potential transcriptional systems that regulate these oscillatory programs across cell types, we used regulatory network analysis to predict transcriptional regulators of pulsatile genes. We find that most cell types, other than pharynx, are likely to deploy the same upstream components. Overall, our results support a model in which shared transcriptional regulators control cell-type-specific oscillatory gene expression, including surprisingly diverse programs for cuticle synthesis.

## RESULTS

### A glial-enriched dataset for analysis of oscillatory gene expression

To analyze global patterns of oscillatory gene expression, we reasoned that it was important to sample not only well-studied cell types such as hypodermis but also cell types like glia for which little or no oscillatory gene expression has been described. We also wished to collect samples at various times in the larval cycle.

To this end, we used *C. elegans* strains expressing GFP specifically in ILso glia or in all glia (*grl-18*pro::GFP or *mir-228*pro::GFP, respectively). We performed fluorescence activated cell sorting (FACS) of dissociated cells to enrich for fluorescent glial cells, using low stringency gating parameters to sample other cell types as well (Fig. 1A). From the resulting cells, we collected single-cell RNA-Seq (scRNA-Seq) data. We collected 10 biological replicates targeting populations in the L2 or L4 larval stages (Fig. 1A, Supp. Table S1), resulting in a total of 24,057 cells. Annotation of the resulting dataset identified ILso (9% of total) and other glial cells (23%), as well as other tissue types including skin (23%), muscle (7%), neurons (9%), and pharynx (15%) (Fig. 1B, C; detailed annotations in Supp. Fig. S1 and Supp. Table S2). Notably, there are only six ILso glia per animal (0.6% of total cells), indicating ∼15-fold enrichment of this cell type in our dataset. Together with recent studies that targeted adult glia (Purice et al. 2025) and all cell types across larval stages (D. Gaidatzis, H. Großhans, personal communication), this dataset provides a powerful resource for examining oscillatory gene expression in glia and other understudied cell types.

**Figure 1.**
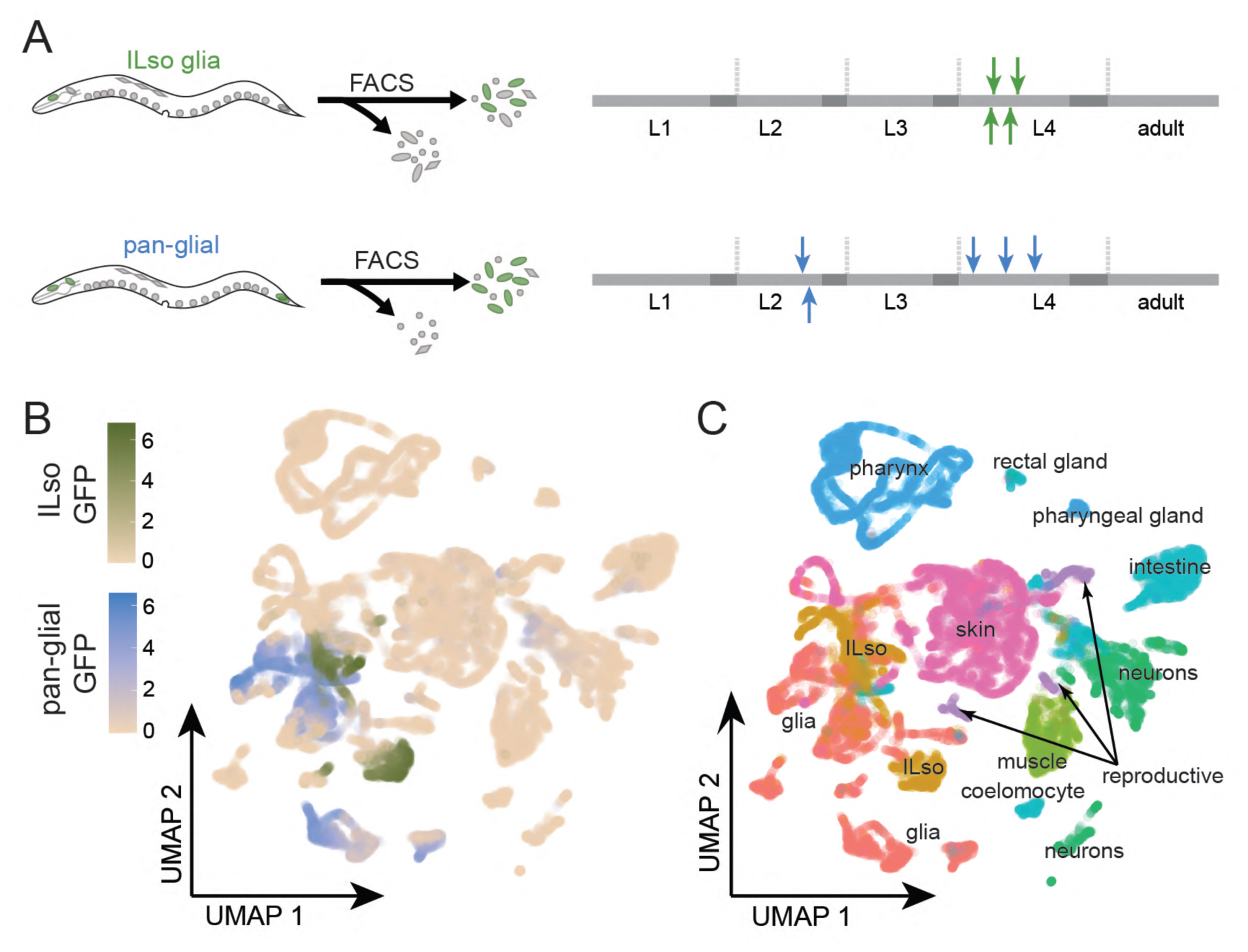
Glia-focused sorts during development reveal cell types that form diverse structures in UMAP space. (A) Sorts targeted glia and ILso for sequencing. Arrows indicate time of bulk animal collection relative to standard worm development; the stage of individual animals is variable. (B) UMAP showing recovered cells and expression of the GFP transgene in ILso-targeting sorts (green) and pan-glial sorts (blue). (C) Tissue-level annotation.

### ILso glia exhibit oscillatory gene expression

We first analyzed the ILso glial cells using principal component analysis (PCA). When projected onto the first two principal components, the ILso glial cells organized themselves in a striking circular pattern (Fig. 2A) reminiscent of PCA signatures associated with cyclic gene expression during the cell cycle (Chervov and Zinovyev 2022). We reasoned that this pattern might correspond to oscillatory gene expression during *C. elegans* larval development that was previously observed through whole-animal (bulk) RNA-Seq (Meeuse et al. 2020). Briefly, in that study, 3,739 genes were found to exhibit oscillatory expression across larval stages, such that the peak expression of each gene occurred rhythmically at a consistent time within each larval stage. To quantify the peak timing for each oscillating gene, its expression was fit with a cosine wave (Meeuse et al. 2020). Then, the timing of its peak expression was described as a phase angle between 0° and 360°. Oscillating genes with similar phase angles have peak expression at similar points in the larval cycle (Meeuse et al. 2020).

**Figure 2.**
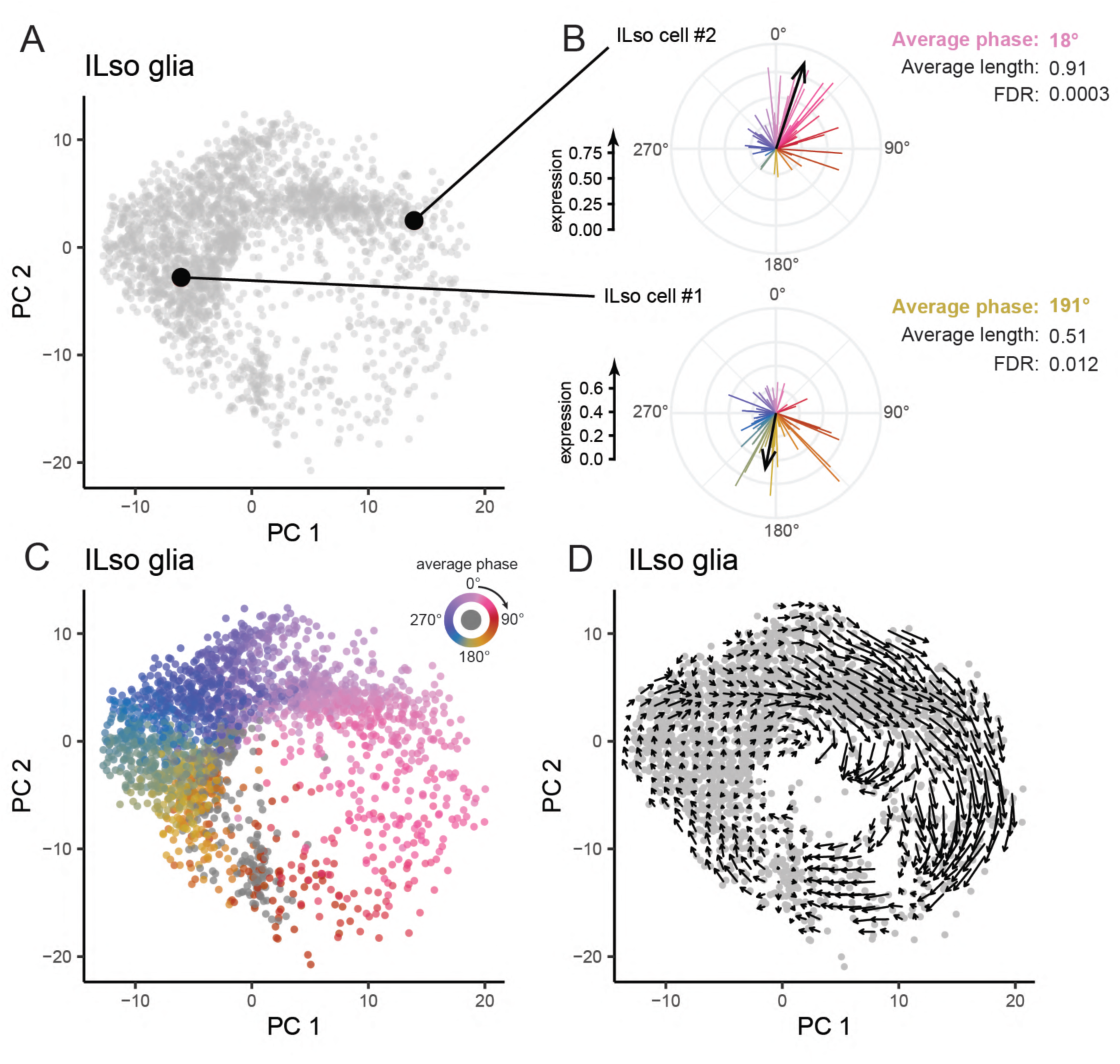
ILso glia exhibit transcriptional oscillations reflecting the larval cycle. (A) PCA plot of ILso glia. Two individual cells are highlighted. (B) Computation of the average phase angle for the two cells highlighted in A. For each cell, the expression level of all genes annotated as oscillating in bulk data (Meeuse et al. 2020) is plotted on a radial graph, where the length of each line indicates the expression in the individual cell, and the angle indicates the oscillation phase from bulk data. The thick arrow indicates the circular weighted average, and colors represent the phase angle (inset). FDR values are calculated using a permutation test. (C) PCA plot of ILso glia, where each cell is colored according to its average phase. Cells colored in grey failed to reject the null hypothesis in the permutation test. (D) RNA velocity superimposed on the PCA plot.

To test if such oscillatory gene expression is present in ILso glia, we computed the average phase of each cell (Fig. 2B). Specifically, for each cell, we computed a weighted circular average of the peak phases of oscillating genes (derived from the previous bulk RNA-Seq data), using the gene expression levels in that cell as weights. This average results in a vector whose direction reflects the average phase of genes expressed in that cell, and whose length reflects how consistently the genes’ peak times align in that cell (Fig. 2B). We found that most ILso glial cells showed a strong average cell phase; that is, the known oscillating genes expressed in each cell had similar peak phases. By contrast, applying the same approach to body wall muscle (BWM) cells, we found that most cells expressed few oscillating genes, with no consistent bias towards a particular phase (Supp. Fig. S2).

To quantify this phenomenon, we permuted the gene phases (Supp. Fig. S2C) and found that 88% of single ILso glial cells, but only 7% of BWM cells, had a statistically significant average phase at FDR < 0.05.

Next, to visualize how the average phases of ILso glia relate to their organization in PCA space, we applied a circular color palette to encode the average phase of each cell (Fig. 2C). Strikingly, this analysis reveals that progression around the circular structure formed by ILso cells in PCA space precisely corresponds to progression through the phase space of cyclic gene expression (Fig. 2C, compare cell color to inset). By contrast, the organization of BWM cells in PCA space does not obviously relate to phase (Supp. Fig. S2D).

As an independent assessment that does not rely on established peak phases, we analyzed RNA velocity in the ILso glia (Fig. 2D). Briefly, while mature mRNA counts are used to define the state of a cell at a fixed timepoint, RNA velocity approaches use the relative abundance of unspliced pre-mRNA to infer the future state of that cell (La Manno et al. 2018). A simple visualization of RNA velocity in PCA space is to draw an arrow from the current state of a cell to the position of its projected future state. Applying this approach to ILso glia, the resulting pattern further suggests that the circular organization of these cells corresponds to progression through a cyclic program of gene expression (Fig. 2D).

Together, these results indicate that the oscillatory patterns previously observed in bulk data can be readily identified at the level of individual cell types where they shape organization of the PCA space, revealing the presence of an oscillatory gene expression program in ILso glia.

### Cuticle-producing cell types display oscillatory gene expression

Next, we sought to apply these approaches to other cell types. As PCA plots are inherently limited in the number of dimensions they can display simultaneously, we used UMAP to visualize many cell types at once (Fig. 3A, B). In each plot, we color-coded cells that had a statistically significant average phase, and represented all other cells in gray. Visualizing average phase in this manner revealed that individual skin, glia, and pharynx cells tend to display a strong average phase that correlates with their organization in UMAP space, while intestine, neurons, and muscle cells do not (Fig. 3A, B). We examined individual cell types using PCA plots color-coded by average phase and overlaid with RNA velocity assessments (Fig. 3C), similar to our analysis of ILso glia. We found that this method further supports the presence of cyclic gene expression programs in specific cell types, for example in seam cells and pharynx epithelial cells, but not in BWM cells or SMB neurons (Fig. 3C).

**Figure 3.**
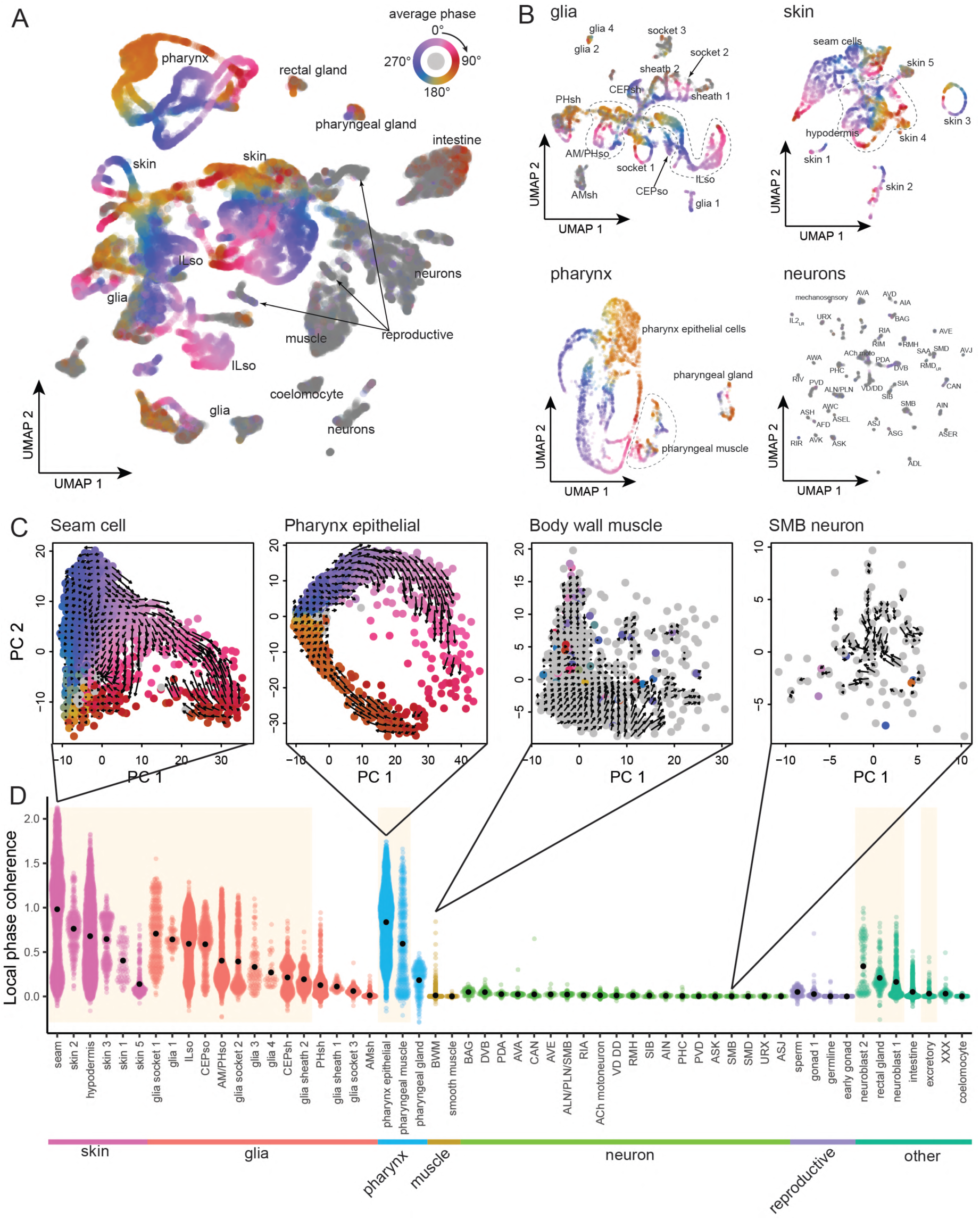
Oscillation patterns across multiple cell types. (A) UMAP of all cells in our dataset colored by average phase. Cells failing the permutation test are colored in grey. (B) UMAPs for selected tissues, indicating individual cell types and colored by average cell phase. Cells failing the permutation test are colored in grey. (C) PCA plot indicating RNA velocity (arrows) and colored by average phase for selected cell types. (D) Beeswarm plot of phase coherence index across all cell types in data set, grouped and colored by tissue. The black dots indicate the mean phase coherence index within each cell type. The colored background highlights the cell types with a statistically significant phase coherence index (p < 0.05 by permutation test).

To more systematically assess differences between cell types, we formulated a metric called local phase coherence that quantifies, for each cell, the similarity of its average phase to that of its 20 nearest neighbor cells in transcriptional space. We used a permutation approach to derive statistical confidence for this metric (Fig. 3D, see Methods). Cell types that have a high local phase coherence are likely to be oscillatory; that is, transcriptional variation within that cell type is strongly explained by cells being caught at different time points in their oscillatory cycle. We found that most skin cell types, glia socket cell types, pharynx epithelial cells, and pharyngeal muscles displayed a high and statistically significant local phase coherence, while neurons, muscle cells, and reproductive cell types did not (Fig. 3D). This indicates that during larval development, the cell types that are involved in cuticle formation (e.g. hypodermis, seam, socket glia, pharynx) show the strongest oscillatory gene expression patterns. By contrast, although neurons and muscles are involved in behavioral changes such as lethargus during the larval cycle, these changes are likely not driven by large-scale cyclic changes in gene transcription.

### *De novo* identification of pulsatile genes

Next, we developed analytical methods to identify pulsatile genes in individual cell types *de novo*.

We reasoned that previous analyses identifying oscillatory genes in whole-animal bulk sequencing might be insensitive to gene expression patterns in rare cell types. Therefore, as a complementary approach, we sought to identify oscillating genes *de novo* using only scRNA-Seq data.

In cell types displaying strong oscillatory patterns, the cyclic pattern of overall gene expression manifests as a circular shape in two-dimensional PCA space. Cell position around the circle defines a pseudotime, with each orbit around the circle corresponding to one full cycle of larval development.

We reasoned that for individual genes, if gene expression in a given cell type were plotted as a function of pseudotime, oscillatory genes would display a distinct peak because they are expressed at a particular pseudotime (Fig. 4A). By contrast, we expected to see flat or complex shapes for genes in the same cell type that are expressed constitutively or that vary in a manner unrelated to larval oscillations (Supp. Fig. S3A).

**Figure 4.**
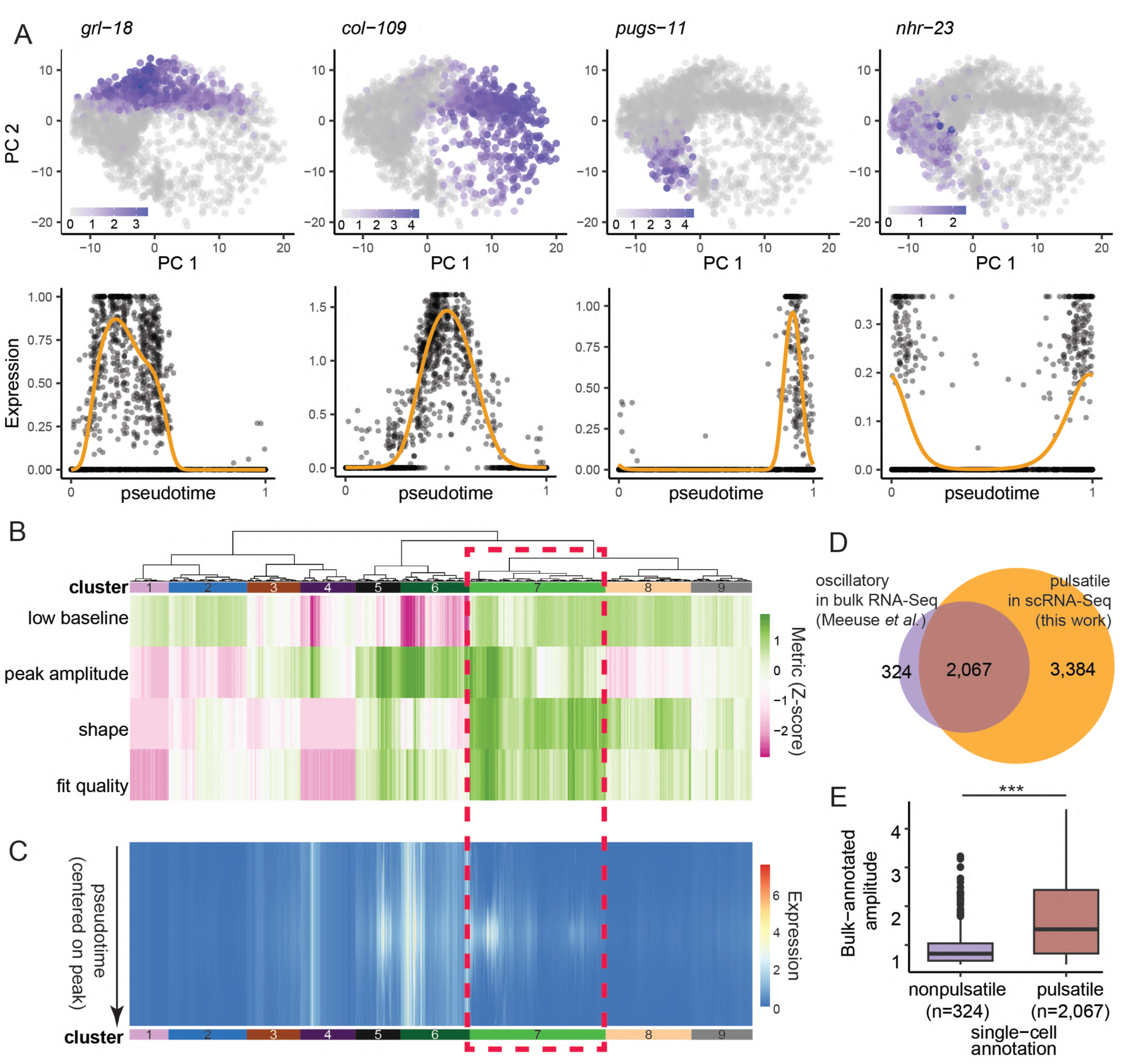
*De novo* identification of pulsatile genes. (A) Single-cell expression levels and GAM fits for 4 genes in ILso. Top row: PCA plots, each cell is colored by the raw number of reads for the indicated gene (log scale). Bottom row: cells are ordered by pseudotime, with normalized gene expression (log scale) on the y axis, clipped at the 95^th^ percentile. The orange line indicates the GAM fit. (B-C) Heatmap of predictors (B) and gene expression (C) across clusters. Red box indicates Cluster 7, containing the genes considered to be pulsatile. (D) Overlap of pulsatile genes in the 22 oscillatory cell types described here with bulk-annotated oscillating genes (Meeuse et al. 2020). (E) Bulk-annotated oscillation amplitude among pulsatile and non-pulsatile genes, only examining previously reported bulk genes (Meeuse et al. 2020). Two-sided Wilcoxon test: p < 1e-16.

Thus, for each oscillatory cell type, we inferred a circular pseudotime by fitting an elastic principal graph in PCA space (Supp. Fig. S3B, see Methods). We then fitted the expression pattern of each gene with a generalized additive model (GAM) to obtain smoothed expression profiles. Next, we developed a quantitative criterion to identify genes that exhibit a well-defined peak in pseudotime.

Briefly, we derived four descriptors from each curve (Supp. Fig. S3C; Supp. Table S3): baseline expression (the background level outside of the pulse), peak amplitude (the maximum expression during the pulse), shape (how closely the curve resembles an ideal sharp peak), and fit quality (how well the GAM captures actual expression along pseudotime). Importantly, each oscillatory cell type was analyzed separately in case a given gene had pulsatile expression in one cell type but not another. Finally, we performed hierarchical clustering to group the gene-cell type curves in an unbiased manner based solely on these descriptors (Fig. 4B, C; Supp. Fig. S3D; Supp. Table S3).

Among the resulting clusters, Cluster 7 genes best match our four criteria: they display a low baseline, high peak expression, peak-like shape, and high fit quality. Thus, we infer that this cluster contains pulsatile genes (red box in Fig. 4B, C). Clusters 8 and 9 contain genes that are potentially pulsatile but with lower peak amplitude (Fig. 4B, C). By contrast, the genes in Clusters 1 and 4 were not accurately captured by the GAM, as evidenced by their low fit quality, while genes in Clusters 3 and 6 had a high baseline, and thus are likely expressed constitutively at the transcript level (Fig. 4B, C).

Altogether, Cluster 7 contained 23,362 curves that we classified as pulsatile, corresponding to 8,653 unique genes (some of which are pulsatile in more than one cell type). Among these, we focused on those genes that were pulsatile within the 22 cell types we identified as oscillatory based on local phase coherence (Fig. 3D), resulting in 15,460 pulsatile curves, corresponding to 5,451 unique genes. A summary of all genes with pulsatile expression in each cell type is in Supp. Table S4. The 5,451 genes we identified as pulsatile in oscillatory cell types using our *de novo* approach include 2,067 of the 2,391 genes (86%) that were previously identified as oscillating by bulk RNA-Seq, as well as an additional 3,384 genes that were not previously identified as oscillating (Fig. 4D). The small portion of genes annotated as oscillating in bulk RNA-Seq but not in our analysis tended to have shown low amplitude oscillations compared to those supported by both approaches and thus may reflect either greater sensitivity or vulnerability to noise of the bulk method (Fig. 4E).

Overall, our *de novo* approach using single-cell data largely supports the genes that were previously identified as oscillatory from bulk sequencing and further expands this set by >2.5-fold, suggesting ∼25% of *C. elegans* genes exhibit oscillatory expression in one or more cell types.

### *De novo* identification of oscillatory cell types

Next, we leveraged our *de novo* analysis of pulsatile genes to independently derive a list of cell types with oscillatory gene expression. There are two motivations for undertaking this analysis. First, our previous approach using local phase coherence (Fig. 3D) might overlook cell types that express genes that were not annotated as oscillatory in the bulk data, but were identified as pulsatile in our single-cell analysis (Fig. 4D). Second, such a computational approach would be valuable in other cases where only broadly-timed single-cell data exist, without availability of bulk data from carefully synchronized time points.

We reasoned that, in an oscillatory cell type such as ILso glia, pulsatile genes are expressed at various points along the pseudotime trajectory, such that gene ensembles appear as a continuous wave of gene expression (ILso glia in Fig. 5A, B). By comparison, in non-oscillatory cell types, PCA structure typically reflects a small number of discrete cell states, with gene ensembles expressed uniformly in distinct regions of the PCA plot rather than as a continuum (body wall muscle in Fig. 5A, B; Supp. Fig. S4). We thus calculated the perplexity (the exponentiated entropy) of peak timings within each cell type (see Methods). High perplexity reflects a relatively uniform (wave-like) distribution of peak timings, whereas low perplexity indicates a biased (step-like) timing of expression peaks. We hypothesized that this metric would distinguish between pulsatile cell types (corresponding to relatively high perplexity) and non-pulsatile cell types (corresponding to low perplexity).

**Figure 5.**
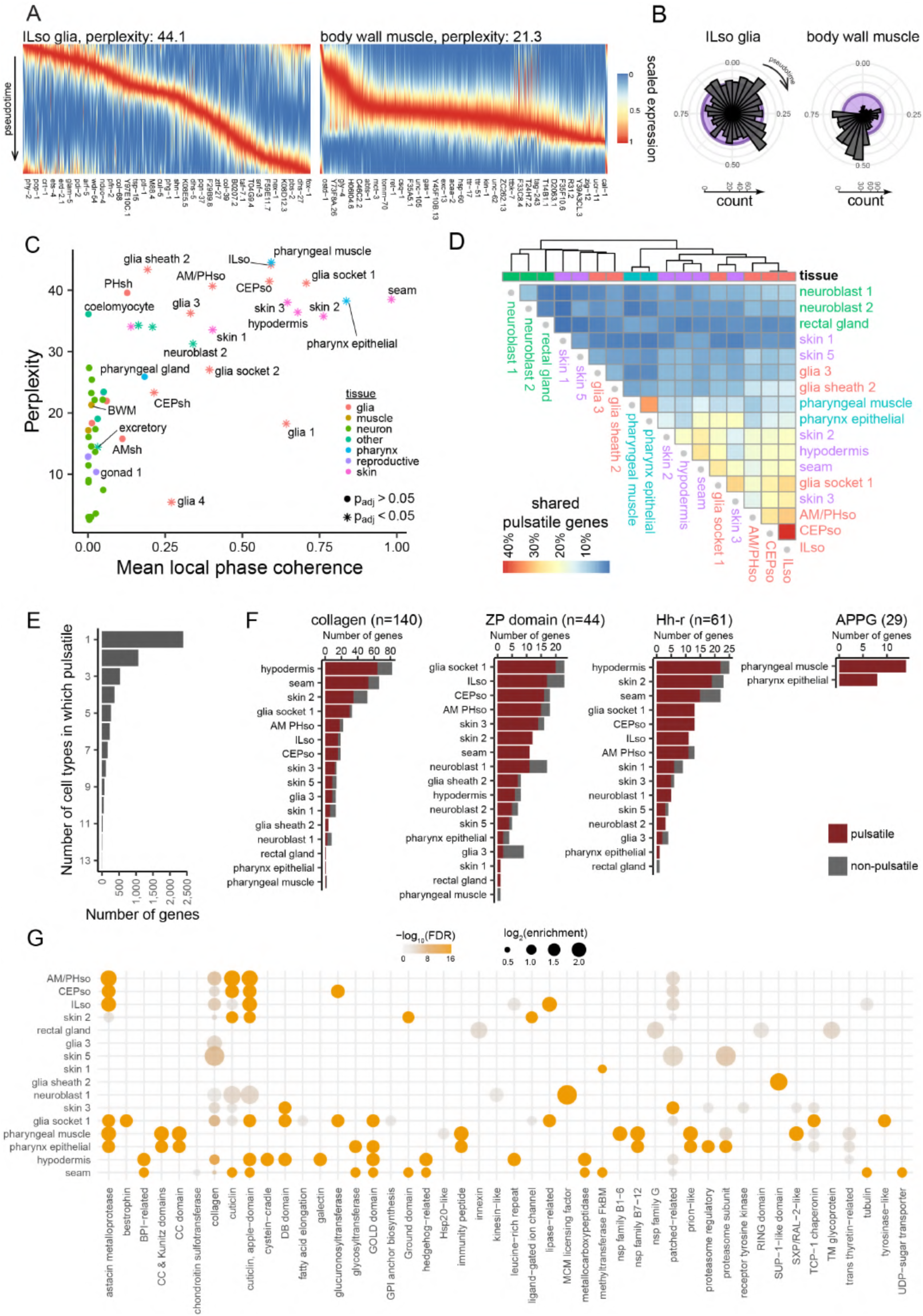
Pulsatile genes across cell types. (A) Expression of the pulsatile genes in ILso glia (left) and body wall muscle (right). The genes are ordered by their time of peak expression. (B) Histogram of the pseudotime (shown as phase) of peak expression of all pulsatile genes in ILso (left) and body wall muscle (right). Each bin = 12 degrees. (C) Perplexity and mean local phase coherence (as described in Fig. 2) for each cell type. Stars indicate a statistically significant (p_adj_ < 0.05) local phase coherence. (D) Overlaps of pulsatile genes in oscillatory cell types. For each pair of oscillatory cell types, the color indicates the proportion of genes pulsatile in both cell types, among genes pulsatile in one of the cell types of the pair. (E) Number of cell types in which each gene is detected as pulsatile. (F) For four selected gene families known to be involved in cuticle production, number of genes expressed and pulsatile in each oscillatory cell type. Gene families: collagens, ZP domain proteins, Hedgehog-related (Hh-r), ABU/PQN Paralog Group (APPG). The number of total genes in each family is indicated in parentheses. (G) PANTHER families enrichment in pulsatile genes for each oscillatory cell type. Dot size indicates enrichment fold change, dot color and transparency indicates statistical significance.

To assess whether higher perplexity corresponds to oscillatory gene programs, we plotted perplexity against local phase coherence for each cell type (Fig. 5C; Supp. Table S5). We find that 17 of the 19 cell types with the highest perplexity (scores >30) are among the 22 cell types we identified using local phase coherence (Fig. 4E, significant cell types represented as stars in Fig. 5C). These include all skin-related cell types as well as many glial and pharyngeal cell types (Fig. 5C). Thus, *de novo* identification of oscillatory cell types using perplexity of pulsatile gene expression largely recapitulates our analysis using bulk-derived phase coherence in PCA space. We propose that perplexity of time of peak expression is a useful tool for *de novo* identification of oscillatory cell types using only scRNA-Seq data.

Two cell types (coelomocytes and PHsh glia) had perplexity > 30 but were not identified as oscillatory using local phase coherence, while four cell types (CEPsh, glia1, glia4, excretory) had statistically significant phase coherence but low perplexity. These non-overlapping cell types likely represent differential strengths and weaknesses of the two approaches (see Discussion). We limit our subsequent analyses to the 17 high-confidence cell types that appear oscillatory using both approaches. A summary of pulsatile genes in these 17 cell types is shown in Table 1.

**Table 1.** Overlap between pulsatile genes from *de novo* analysis of scRNA-Seq and genes previously identified as oscillatory in bulk RNA sequencing.

|  |  | **single-cell** |  |  |
| --- | --- | --- | --- | --- |
|  |  | non-pulsatile | pulsatile |  |
| **bulk** | non-oscillating | 2,792 | 3,236 | 6,028 |
|  | oscillating | 335 | 2,032 | 2,367 |
|  |  | 3,127 | 5,268 | 8,395 |

### Pulsatile genes differ across tissues and are involved in cuticle formation

To what extent are the same sets of pulsatile genes shared between cell types? To address this question, we examined the overlap between the pulsatile genes we identified in each cell type (Fig. 5D). We found that cell types belonging to the same tissue display more similarity, for example pulsatile genes are more likely to be shared within glial cell types (ILso, CEPso and AM/PHso), skin cell types (seam cells, hypodermis, and the unidentified skin 2 cluster), and pharynx cell types (pharyngeal muscle and pharyngeal epithelia). However, this overlap represents at most 40% of the pulsatile genes (in the case of ILso and CEPso), with many cell types sharing <10% of their pulsatile genes (Fig. 5D). Put another way, 45% (2,390 of 5,268) of the genes we identified were expressed and pulsatile exclusively in a single cell type while only 17% (915 of 5,268) were pulsatile in five or more cell types (Fig. 5E). Interestingly, only 10 genes were pulsatile in all 17 oscillatory cell types, and these include *lin-42/*Period and *myrf-1/*MYRF, which have been implicated as critical timing regulators in larval development (McCulloch and Rougvie 2014; Van Wynsberghe and Pasquinelli 2014; Edelman et al. 2016; Meeuse et al. 2023; Spangler et al. 2024; Xu et al. 2024). Together, this suggests that different cell types use largely distinct repertoires of oscillating genes.

We investigated the identity of protein families encoded by genes with pulsatile expression. We initially focused on four gene families known to be involved in patterning the cuticle and precuticle (a transient aECM thought to scaffold patterning of the mature cuticle (Sundaram and Pujol 2024)): collagens and ZP domain proteins/cuticlins, which are two of the major structural components of cuticle (Lazetic and Fay 2017; Sundaram and Pujol 2024); Hedgehog-related (Hh-r) proteins, which have recently been shown to be involved in cuticle patterning (Fung et al. 2023; Serra et al. 2024); and ABU/PQN paralog group (APPG) genes, which are involved in formation of the pharyngeal cuticle (George-Raizen et al. 2014). We found that most collagens, cuticlins, and Hh-r genes are pulsatile in at least one cell type among the skin and socket glia, consistent with a major role for these gene families in larval development (Fig. 5F; Supp. Table S4). For example, of the 140 collagen genes, 119 were expressed in at least one cell type, and 107 of these showed pulsatile expression (Fig. 5F; Supp. Table S4). However, these gene families are largely not expressed in the pharynx (Fig. 5F; Supp. Table S4). Conversely, the APPG genes are expressed and pulsatile in the pharyngeal muscles and pharyngeal epithelium, but are not expressed in the skin or glia (Fig. 5F; Supp. Table S4).

We then aimed to extend this analysis by using an unbiased approach to systematically uncover gene families that are particularly prone to displaying oscillations. We downloaded the PANTHER ontology database, which defines gene families based on orthology of the protein sequences (Thomas et al. 2022). We reasoned that this classification is more relevant in our case than ontologies based on gene function or pathway due to the limited functional annotation for genes involved in cuticle formation. We found 47 gene families with significant enrichment in pulsatile genes, including collagens, cuticlins, Hh-r genes and several other families associated with cuticle formation (Fig. 5G, Supp. Table S6). These results support our previous targeted analysis of collagens, ZP-domain, Hh-r, and APPG families, all of which were independently rediscovered by this unbiased approach. In addition, the nematode astacin metalloproteases appeared enriched in pulsatile genes in glia and pharynx, but not hypodermis (Fig. 5G, Supp. Table S6). Conversely, galectins, cysteine-cradle, and GOLD-domain gene families were pulsatile in the hypodermis, but not glia (Fig. 5G, Supp. Table S6). Several families of peptides with putative roles in immunity and stress response showed enrichment specifically in pharyngeal cell types (Fig. 5G, Supp. Table S6).

Taken together, our results suggest that cuticle formation is the main commonality among pulsatile genes, and that distinct cell types use very different gene expression programs during this process. Thus, while cuticle aECM is typically perceived as a single homogeneous meshwork, our results suggest that the cuticle is actually a patchwork matrix with different patterning and composition contributed by distinct cell types.

### Transcription factors driving oscillations in each cell type

Finally, we sought to identify transcription factors (TFs) that may drive expression of the pulsatile genes in each cell type. We used the previously established gene regulatory network CelEst (Perez 2025), which integrates three types of evidence – the presence of TF binding motifs, ChIP-Seq associations, and yeast one-hybrid assays – to predict likely targets of each TF. Using these predicted targets as input, we performed cell-type-specific tests to identify TFs whose targets are statistically enriched in pulsatile genes.

For most cell types in our dataset, we found TFs that were significantly associated with pulsatile genes (Fig. 6A; Supp. Fig. S5A). Notably, *blmp-1*, *nhr-23*, *nhr-25, nhr-41* and *nhr-85* display a significant enrichment in pulsatile genes in their targets in the skin and glial types, consistent with previous evidence suggesting their role in larval oscillations and molting (Johnson et al. 2023; Kinney et al. 2023; Meeuse et al. 2023) (Supp. Table S7). This is consistent with the model that, although different pulsatile genes are expressed in each cell type, they are controlled by shared upstream TFs. By contrast, the pharyngeal cell types did not display an enrichment in the targets of these genes, showing instead 23 TFs, including *bcl-11*, *klu-2* and *nhr-142,* suggesting the pharynx may use an oscillatory mechanism different from skin and glia.

**Figure 6.**
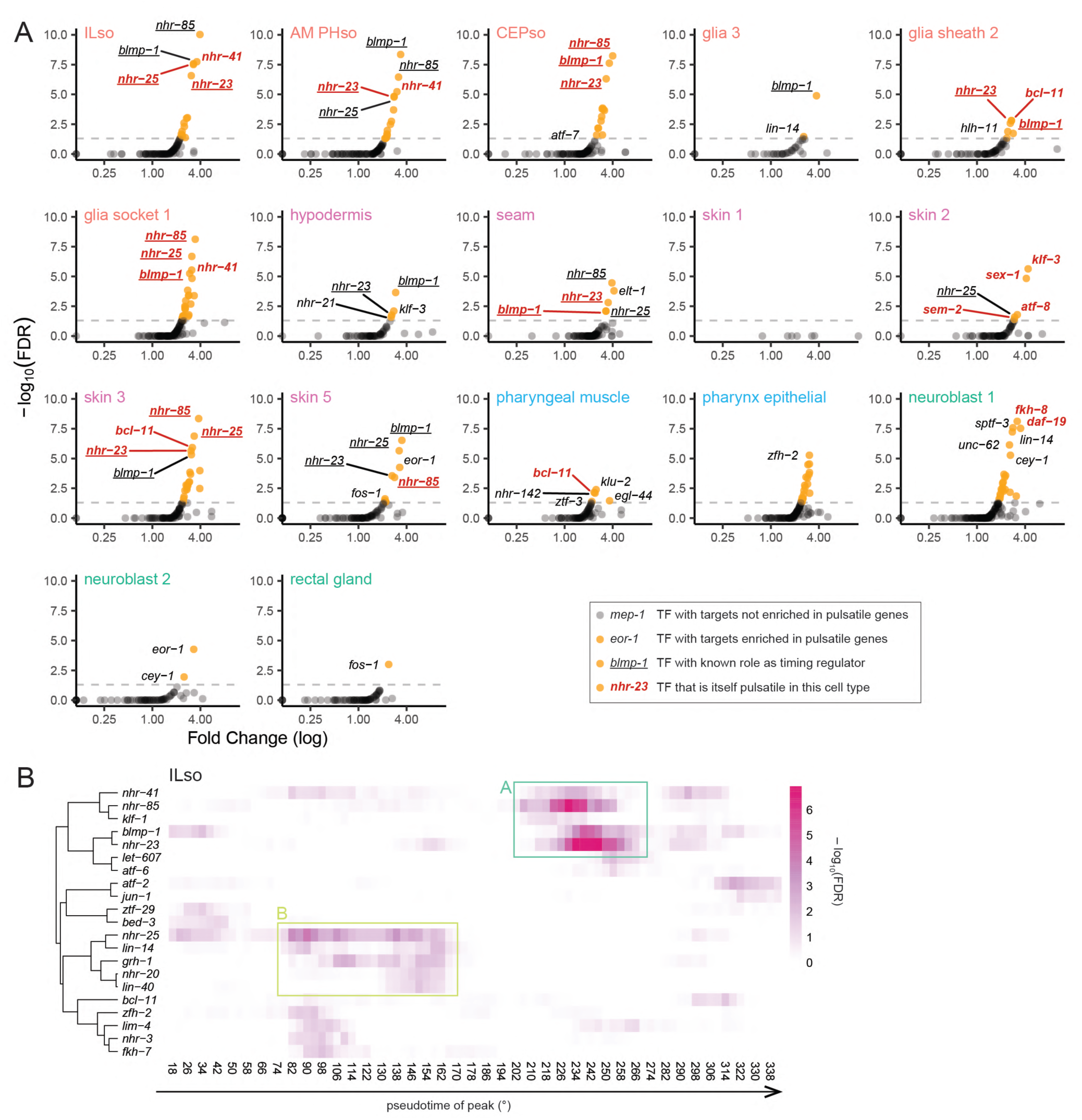
Transcription factors associated with pulsatile genes. (A) Volcano plots indicating, for each oscillatory cell type, the transcription factors (TFs) whose targets are enriched in pulsatile genes. TFs that are themselves pulsatile in this cell type are indicated in bold red, TFs with a reported role in regulation of oscillations are underlined. (B) TFs with targets enriched in genes pulsating in ILso and peaking within a time bin. Overlapping time bins are represented along the horizontal axis and mapped to the bulk phases. The color indicates statistical significance. The boxes highlight two groups of interest.

We reasoned that if these TFs are driving oscillatory expression, they may act at specific phases in each larval stage. To test this hypothesis, we binned the pulsatile genes based on their time of peak expression and tested TFs for significant association with pulsatile genes peaking around a particular phase (see Methods). In ILso glia, we observed two groups of TFs whose targets were enriched in genes peaking at particular times. Group A contained *blmp-1*, *nhr-23*, *nhr-41* and *nhr-85* and was associated with genes peaking around 240° (Fig. 6B). Group B contained *grh-1*, *lin-14* and *nhr-25*, and was associated with genes peaking around 130° (Fig. 6B). Repeating the analysis in other cell types, groups A and B could be observed at similar timepoints in the other skin and glial cell types (Supp. Fig. S5B). In pharyngeal cell types, we did not observe enrichment of these two groups, but instead we identified group C, containing *pha-4* and associated with genes peaking around 100° and group D, containing *let-607* and associated with genes peaking around 270° (Supp. Fig. S5B).

Together, this suggests that shared groups of TFs – an “early” and “late” group in skin and glia, and a different early and late group in pharynx – may drive expression of cell-type-specific gene sets at particular times during the larval molt cycle.

## DISCUSSION

### Analytical tools to uncover oscillatory patterns in single-cell transcriptomics

Oscillatory gene expression occurs across biological systems – including the cell cycle and circadian rhythms, making *de novo* identification of oscillatory genes a key biological problem. While several computational methods have been developed to address this task (Chervov and Zinovyev, 2022), these tools were poorly suited to our context. Cell cycle algorithms rely on known oscillatory gene sets, and methods for circadian or somitogenesis rhythms often assume slower cycles measured with bulk methods. Our work introduces an alternative approach that leverages the power of single-cell RNA sequencing to discover oscillatory genes. In principle, our approach can be applied to any scRNA-seq dataset where oscillations dominate the PCA space. We expect these methods to be useful for identifying oscillatory genes in other contexts.

Because our method exploits PCA structure, it works effectively when oscillations dominate the principal component space. This is nicely illustrated in cuticle-forming cells, where the first two components form a circular trajectory corresponding to pseudotime. By contrast, other cell types, such as muscle or neurons, do not show this pattern. This could indicate either a genuine absence of oscillatory programs or the presence of oscillations driven by only a few genes that are insufficient to shape PCA structure. Notably, the pulsatile microRNA *lin-4* cycles in intestinal cells (Kinney, 2024), and neuronal oscillations – likely linked to sleep during lethargus – have been reported (Turek and Bringmann, 2014), suggesting that limited gene sets can still drive physiologically relevant rhythms.

### Socket glia express cell-type-specific cuticle aECM genes

While prior work has focused on oscillations in the hypodermis, our dataset extends the analysis to diverse tissues, including glia. We captured a broad range of cell types (Supp. Table S2) and enriched for ILso and other glial populations through targeted sorting. ILso cells – rare in abundance at only six cells per animal – have been difficult to identify in previous single-cell datasets (Cao, Packer et al. 2017; Taylor, Santpere et al. 2021). Their unambiguous detection here underscores the sensitivity of our approach and complements recent advances in glial cell profiling (Purice, 2025).

Previous work showed that some of the most highly subtype-specific glial markers are genes that encode aECM proteins (e.g, *grl-2* in AM/PHso; *grl-18, col-53,* and *col-177* in ILso (Fung et al. 2020)). Consistent with this idea, we find that socket glia exhibit strong oscillatory gene expression throughout the larval cycle, including dozens of putative cuticle components such as collagens, ZP domain proteins, and Hh-r proteins. The observation that socket glial cells cycle through multiple transcriptional states may partly explain why each glial type may appear to populate multiple clusters in some analyses (Purice et al. 2025). Importantly, we find that differential expression of oscillatory aECM genes is one of the main features that distinguishes socket glial cells belonging to different sense organs. This suggests that a defining role of socket glia is to produce cuticle aECM specialized for the function of the sensory neurons they support.

### Spatial and temporal patterning of cuticle aECM

Far from being a simple meshwork, cuticle aECM has been shown to exhibit extensive spatial patterning. Cuticle patterning is achieved in part by localized subcellular assembly of components (e.g., struts, alae, or furrows in the main body cuticle) but also through cell-specific differences in gene expression. For example, the ZP domain proteins DYF-7 and CUTL-18 are exclusively expressed by and localize to subsets of tube-shaped epithelia (sense organs, excretory duct/pore, vulva, rectum) (Heiman and Shaham 2009; Low et al. 2019; Schmidt et al. 2025). Further, high-throughput tagging of >100 endogenous aECM proteins revealed extensive tissue-specific localization patterns (Ragle et al. 2025). One interpretation is that a shared “generic” cuticle becomes modified with various cell-type-specific decorations. However, our results suggest that the bulk of the cuticle itself is likely to be divergent across tissues, typically with <20% of components shared between cell types. aECM composition may thus be seen as a marker of cell identity for many epithelial cells, much as neurotransmitter type is for neurons. The functional roles for such cell-specific differences in the cuticle remain to be explored.

The cuticle also exhibits temporal patterning. At each molt, a transient aECM called the precuticle precedes formation of the mature cuticle (Sundaram and Pujol 2024). Some families of aECM proteins have been proposed to contribute differentially to each structure, for example nidogen domain proteins in precuticle and collagens in mature cuticle (Sundaram and Pujol 2024). Consistent with the concept of early and late genes in cuticle synthesis, we identified early and late groups of TFs based on the phase of their targets. These TFs are shared among skin and glial cell types, but differ in pharyngeal cell types. It is tempting to speculate that the early group, consisting of *blmp-1*, *nhr-23*, *nhr-41* and *nhr-85* in skin and glia, might regulate precuticle genes, while the late group, consisting of *grh-1*, *lin-14* and *nhr-25* in these cells, might regulate mature cuticle genes. However, we were unable to detect enrichment of specific gene families, such as nidogen domain proteins, among the early or late gene targets. It is possible that gene families do not map neatly to precuticle or cuticle, for example GRL-18 is only in precuticle in sense organs but remains in the mature cuticle in the vulva (Fung et al. 2023; Serra et al. 2024). However, it is also likely that post-transcriptional mechanisms not captured in our data contribute to the timing of precuticle and cuticle gene expression.

### Limitations of the study

Because our dataset focused primarily on L4 animals, with more limited representation from L2, we may have missed stage-specific differences. For example, the main body cuticle contains alae only in L1, dauer, and adults, presumably corresponding to differences in gene expression at these stages (Katz et al. 2022). Further, as a result of our sampling methodology, temporal dynamics during L4 cannot be readily distinguished from *bona fide* multi-stage oscillations and genes expressed only in specific regions of PCA space due to tissue heterogeneity (e.g. anterior vs posterior body wall muscle) may appear pulsatile without truly oscillating across development. Broader temporal sampling across all larval stages (including dauers) would mitigate these issues and shed light on the interesting question of how stage-specific differences arise.

The complete transcriptional network that guides oscillating gene expression across all tissues remains unknown. Our analysis identified strong candidates for transcription factors that regulate sets of pulsatile genes. But likely additional factors also play a role, and the question of how these transcription factors are themselves regulated is not addressed by our analysis.

Finally, our study is restricted to transcript-level measurements of mRNAs. Whether these mRNA oscillations are translated into corresponding protein rhythms remains to be determined, though prior work suggests that oscillatory translation and regulated protein stability could contribute (Hendricks, 2014). Conversely, protein levels may oscillate in physiologically meaningful ways through post-transcriptional mechanisms, such as regulated translation or protein degradation, that are not reflected in transcript levels. Other RNA species such as microRNAs are known to play key roles in developmental timing, but were not collected in our study. Addressing these gaps will require integration of additional measurements with single-cell transcriptomics.

## METHODS

### Data collection and processing

Samples were collected as described elsewhere (Taylor et al. 2024). Briefly, a population of gravid animals was submitted to hypochlorite treatment to isolate eggs. The embryos were allowed to hatch in M9 buffer overnight and starved L1 larvae were transferred to 150 mm diameter plates of 8P peptone-rich growth medium seeded with NA22 bacteria. Animals were then grown at 20°C on food for a given time (Supp. Table S1), then collected by washing the plates with cold M9. The cuticle was weakened with SDS/DTT treatment, and the cells dissociated with pronase E treatment and mechanical trituration with a P200 pipette. The dissociation reaction was monitored under a fluorescent microscope and stopped when a majority of fluorescent cells had been released from the animal bodies. Cells were resuspended in L15 medium with 10% FBS, washed twice, filtered, and sorted with Propidium Iodide or DAPI as viability marker. Cells were counted on a hemocytometer and processed with a 10x Genomics 3’ v3 kit and sequenced with an Illumina Novaseq instrument.

The sequencing data was then processed with CellRanger version 4.0.12, aligning to the WS295 annotation provided by Wormbase (Sternberg et al. 2024) augmented with the addition of a GFP transgene sequence. Ambient RNA was subtracted using SoupX (Young and Behjati 2020), counts normalized with scTransform (Hafemeister and Satija 2019) and cells clustered and annotated with Seurat v 5.2.1 (Hao et al. 2024) using markers from published scRNA-Seq data (Taylor et al. 2021). Briefly, samples from the same stage (L2 or L4) were merged and clustered into individual tissue types. Tissues were then individually reclustered into individual cell types (cell type descriptions and marker genes in Supp. Table S2). The cells corresponding to the same cell type at different stages were then merged for subsequent analysis.

### Average cell phase

We used the table provided in (Meeuse et al. 2020), filtered to consider only genes with OscAmplitude above 1.5. From our single cell data, we used the scTransform-normalized read counts and further scaled each gene by its maximum expression across all cells.

The average angle for cell *𝑖*is then computed as

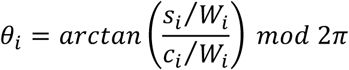

And average length as

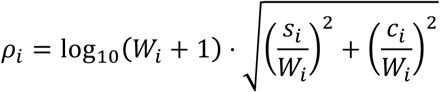

Where 𝜙_*j*_ is the bulk-annotated peak phase (Meeuse et al. 2020), 𝑤_*ij*_ the normalized expression of gene 𝑗 in cell 𝑖 and

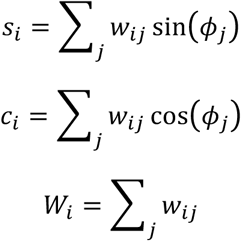

For the permutation tests, we performed 10,000 random permutations of the bulk-annotated phase of peak expression and recomputed the average phase length under this null hypothesis. We compare the observed average phase length to the distribution of average phase lengths under permutation to obtain a p-value, and computed the corresponding False Discovery Rate (FDR) across all cells with the Benjamini-Hochberg method. In general, cells with a FDR above 0.05 are represented in grey; cells with a statistically significant average phase length are color-coded with Peter Kovesi’s cyclic color map (Kovesi 2015).

### Local Phase Coherence index

For each cell, we selected its 20 nearest neighbors calculated with Seurat (Hao et al. 2024). We then computed the dot products of its cell phase with the cell phase of its neighbors. The mean across neighbors was then used as Local Phase Coherence index. For a cell *𝑖* with nearest neighbors *𝑁*(*𝑖*):

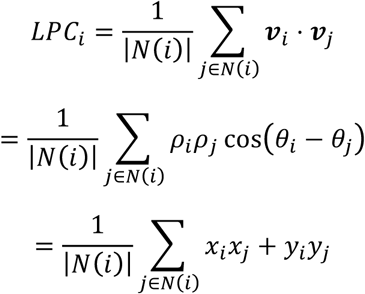

With 𝒗*_i_* = (𝑥*_i_*, 𝑦*_j_*) = 𝜌*_i_* (cos 𝜃*_i_*, sin 𝜃*_j_*).

To further test for statistically significant local coherence, we performed a permutation test, computing the mean Local Phase Coherence index of each cell type after random permutation of the bulk-annotated phase of peak expression. We used 10,000 replications to compute a p-value, that we adjusted for multiple comparison with Holm’s method.

### Gene fitting along pseudotime

The cells were grouped by cell type independent of the stage of collection (L2 or L4), and each cell type was processed individually. Only cell types represented by at least 25 cells were considered.

The SoupX-corrected, unnormalized, counts were used for k-nearest neighbors smoothing (Wagner et al. 2018) version 2, using k=5 neighbors. Following normalization (Hafemeister and Satija 2019), the first two principal components were computed, and an Elastic Principal Cycle was fitted to define a cell type-specific pseudotime (Saelens et al. 2019; Albergante et al. 2020) using 50 bootstrap replicates and 20 node points. This pseudotime ranges from 0 to 1; the point chosen as origin (pseudotime 0) and the direction of increase are arbitrary. As the pseudotime corresponds to the position on a circle, pseudotime values of 0 or 1 are equivalent.

For each cell type, we defined an “expressed gene” as a gene with non-zero expression in at least 20 cells, and in at least 5 % of the cells. We discarded the cell types with fewer than 5 expressed genes. The expression (unsmoothed, scTransform-normalized) of each expressed gene was then fit with a Generalized Additive Model (GAM) using the mgcv package (Wood 2011), with a negative binomial family and a log link, using cyclic cubic regression splines with a basis of size 6. We further included a cell-specific offset using normalization factors computed with the edgeR package (Robinson et al. 2010).

We used the trained model to predict expression of each gene along a grid of 128 regularly spaced pseudotime values, resulting in a smoothed expression profile. Further, for each gene, we shifted the pseudotime values to center the maximal expression value, and fitted a GAM as described above. This yielded a centered, smoothed expression profile for each gene.

### Pulsatile gene identification

For each gene and cell type, we computed four predictors (Fig. S3C). From the centered smoothed expression profile, we used the mean of the 20% lowest values as *baseline expression*. We then computed the exponential of the negative log to obtain a *low baseline* metric where a higher value reflects a lower baseline. The maximum value of the curve was log-transformed and winsorized at the 5%-quantile to be used as the *peak amplitude*. We then scaled the curve by its maximum value, and created an ideal sharp peak as the density of a normal distribution of mean 0.5 and standard deviation 0.01. The Dynamic Time Warp distance between the scaled expression and the ideal peak was computed with the dtw package (Giorgino 2009). We transformed this distance with an exponential decay rate of 0.05 to obtain the *shape* descriptor. Finally, we used the deviance explained (computed by mgcv (Wood 2011)) as a measure of *fit quality*. All descriptors were normalized with a Box-Cox transformation and scaled into Z-scores.

We also considered additional metrics (available in Supp. Table S3) and processing steps, but they did not improve clustering quality. Hence, we restricted our final analysis to these four easily interpretable metrics. All genes and cell types were considered together and submitted to hierarchical clustering based on Euclidean distance and Ward.D2 method, using the fastcluster implementation (Müllner 2013). The resulting clusters were then annotated and an appropriate number of clusters chosen by visual exploration.

### Perplexity

For each pulsatile gene, we define its time of peak expression as the pseudo-timepoint at which the smoothed fit reaches its maximum value. For each cell type, we used 50 bins and obtained the counts of the number of pulsatile genes peaking within each bin. We computed the entropy using the default maximum likelihood method in the entropy R package (Hausser and Strimmer 2025).

Perplexity was obtained by taking the exponential of the entropy.

### Enrichment of gene families

For manually curated gene families, we recovered 61 Hedgehog-related genes compiled in (Hao et al. 2006), and 29 ABU/PQN Paralog Group genes compiled in (George-Raizen et al. 2014). In addition, we downloaded from InterPro 38 ZP-domain genes listed with accession number IPR001507, and 140 collagen genes with accession IPR008160.

We used the Bioconductor package PANTHER.db (Muller 2023) to interrogate the PANTHER database (Thomas et al. 2022) version 18.0 (source date 2023-Sep20). We recovered the gene family annotations for *C. elegans*, containing 14,789 families for 14,472 genes. We restricted our analysis to the 686 families containing more than 5 genes. We then performed a hypergeometric test for each family and each cell type, testing for enrichment of pulsatile genes compared to a background list of genes expressed in this cell type. We selected the gene families containing at least 5 pulsatile genes, with a FDR below 0.1.

### Transcription factors

We downloaded CelEst v1.1 from the author’s Github repository (Perez 2025). CelEst includes three types of evidence for regulatory interactions: TF binding motif, ChIP-Seq, or Y1H; we kept any reported interaction. For each cell type and TF, we performed a one-sided Fisher exact test for enrichment of TF target genes that are pulsatile and computed the FDR with the Benjamini-Hochberg method.

To compare the time of action of the TFs, for each pulsatile gene and cell type, we aligned the pseudotime of peak expression to the bulk-annotated phase by minimizing the error upon shifting the peaks in one degree increments. We then created 90 overlapping bins of width 36 degrees and performed a one-sided Fisher exact test for enrichment of target genes peaking within each bin, followed by FDR computation.

## ACKNOWLEDGMENTS

We thank Oliver Hobert, Robert Aguilar, Abhishek Bhattacharya, Caecilia Thuermer, Wendy Fung, Karolina Mizeracka, Leland Wexler, and Rachel Swope for analysis of glial gene expression, and Aakanksha Singhvi and Helge Grosshans for sharing unpublished work. We thank Keck Microarray Shared Resource (KMSR) at Yale University for assistance with 10x library preparation. Sequencing was performed by the Yale Center for Genomic Analyses at Yale University with support from the National Institute of General Medical Sciences of the National Institutes of Health under Award Number 1S10OD030363-01A1.

