## Supplemental Material for "Systematic identification of oscillatory gene expression in single cell types"

### SUPPORTING TABLES

**Table S1. List of samples** (Format: .XLS)

Summary of strains and conditions used for sample collection; related to Fig. 1A.

**Table S2. Cell type annotations** (Format: .XLS)

Annotation of transcriptional clusters with putative identity, key markers, and larval stage of origin; related to Fig. 1C and Supp. Fig. S1.

**Table S3. Cluster identity of curves (cell x gene)** (Format: .XLS)

Results of hierarchical clustering of gene expression curves in each cell type with descriptors used to characterize peak shape and identify pulsatile genes; related to Fig. 4B.

**Table S4. Gene dynamics and expression across cell types** (Format: .XLS)

Summary of pulsatile or non-pulsatile expression of each gene (n=13,088) in each cell type (n=61); related to Fig. 4 and 5.

**Table S5. Cell type oscillatory criteria** (Format: .XLS)

Perplexity and mean local phase coherence for each cell type, as well as total number of expressed and pulsatile genes detected; related to Fig. 5.

**Table S6. Manual annotation of PANTHER families** (Format: .XLS)

Relevant PANTHER gene family identifiers (PTHR ID) with manually curated summaries of their molecular identity and representative genes; related to Fig. 5G.

**Table S7. Transcription factor enrichment** (Format: .XLS)

Enrichment scores and p-values for each transcription factor's targets among all pulsatile genes in each cell type; related to Fig. 6.

### SUPPORTING FIGURE S1

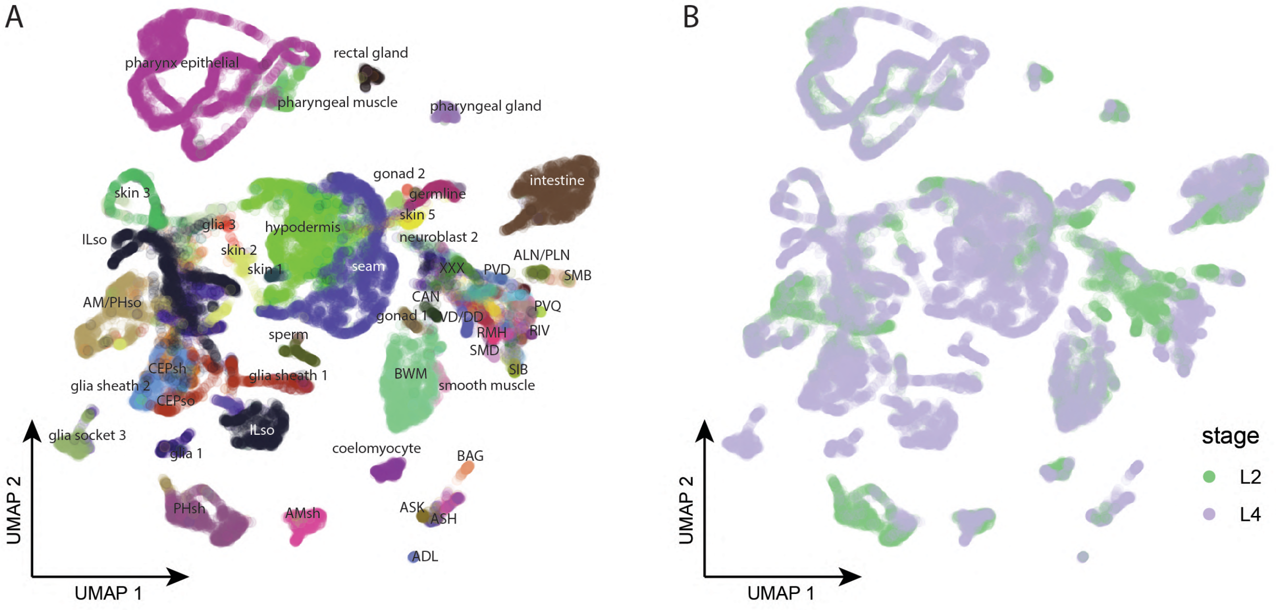


**Supporting Figure S1: Characteristics of the dataset.**

(A) UMAP with selected cell types annotated. (B) UMAP colored by the estimated stage of the animals.

### SUPPORTING FIGURE S2

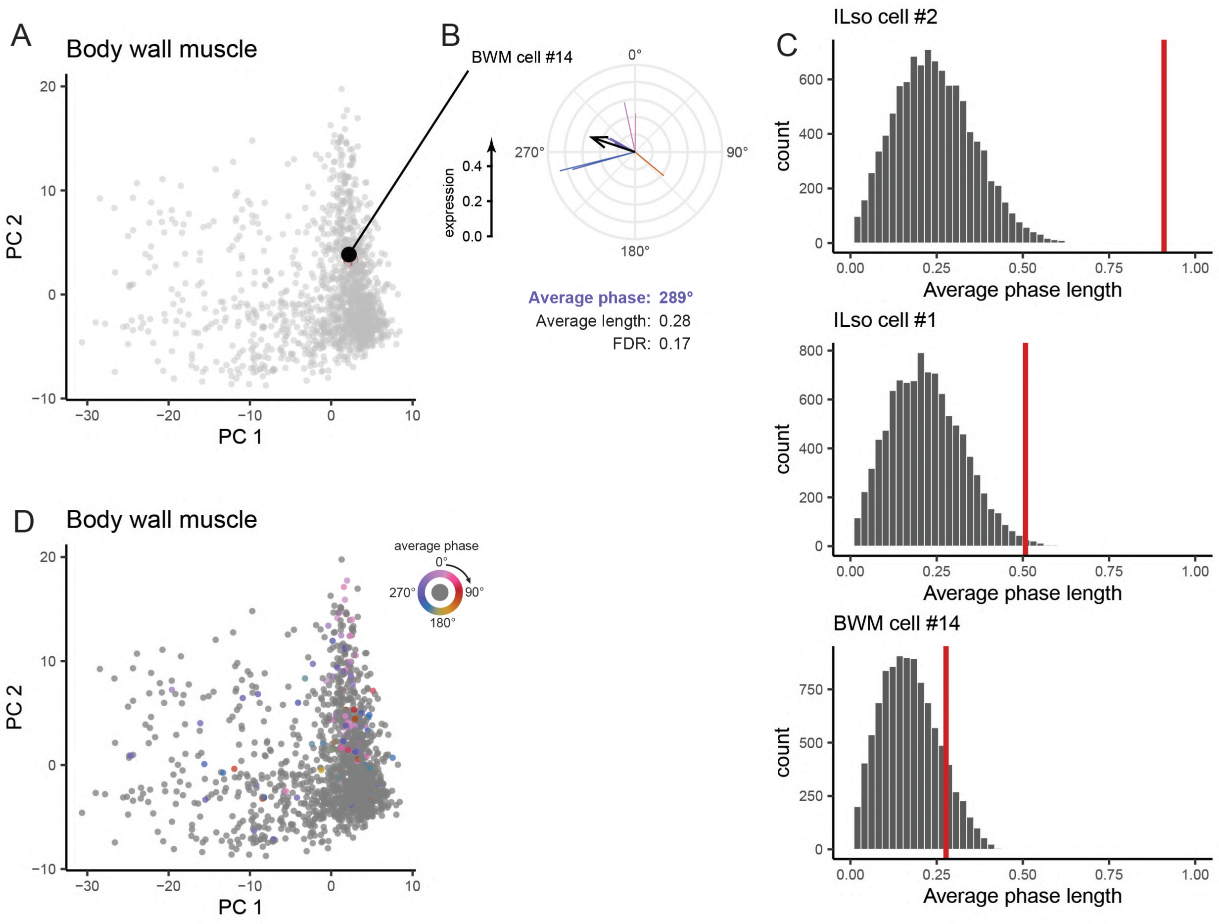


**Supporting Figure S2. Permutation test on average cell phase.**

(A) PCA plot of the body-wall muscle (BWM) cells, highlighting an individual cell. (B) Average phase of the cell highlighted in A. (C) Histogram of the average phase lengths obtained after permutation for the single cells highlighted in ILso and BWM. The red line indicates the (non-permuted) average phase length. (D) PCA plot of the BWM, colored by average cell phase. Cells colored in gray failed to reject the null hypothesis in the permutation test.

**SUPPORTING FIGURE S3**

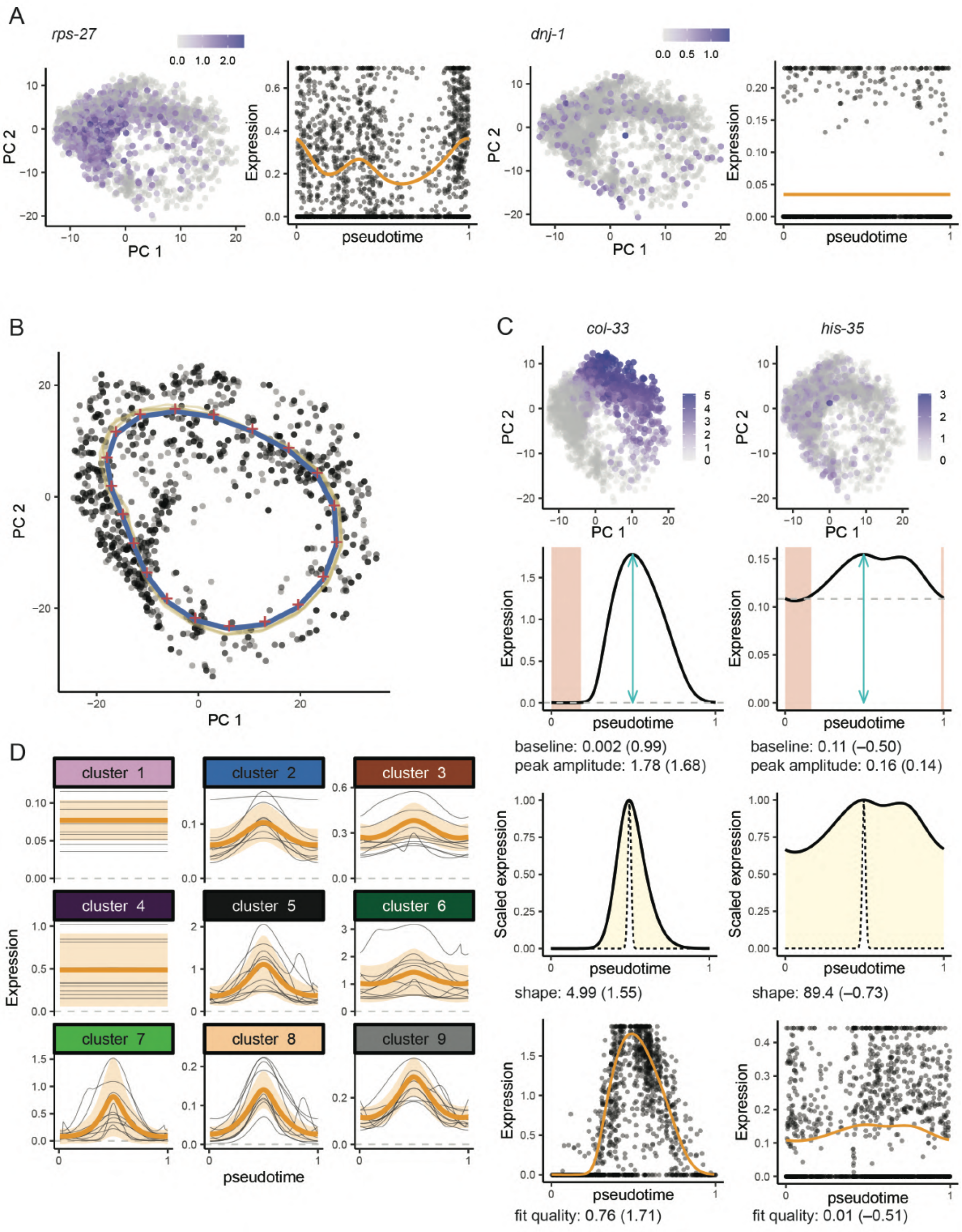


**Supporting Figure S3. GAM fitting approach.**

(A) Example non-pulsatile genes in ILso. Left: PCA plot colored by rawnumber of reads, right: GAM fit on normalized gene expression. (B) Elastic path in the PCA space for ILso. Black dots: single cells, red crosses: node points, yellow lines: 5 bootstrap replicates, blue line: pseudotime trajectory. Origin and direction are arbitrary chosen to define pseudotime. (C) Clustering predictors represented for the genes *col-33* and *his-35* in ILso. The raw value (and Z-score) of each predictor is indicated below the corresponding plot. Top, single-cell expression levels of each gene in ILso. Middle-top, smoothed expression from the centered GAM fit, the peak amplitude is the maximum value of the smoothed expression (cyan arrow), the baseline (dashed gray line) is taken as the mean of the 20% lowest values (highlighted in red). Middle-bottom: scaled smoothed expression (solid line) and ideal sharp peak (dotted line). The Dynamic Time Warping (DTW) distance is symbolized by the colored area. Bottom, the deviance explained by the GAM fit reflects how accurately the expression level of a particular cell (black dots) can be explained by the fit (orange line). (D) Mean expression (orange line) and standard deviation (orange ribbon) within each cluster. For each cluster, 10 randomly selected genes are also plotted (black lines).

### SUPPORTING FIGURE S4

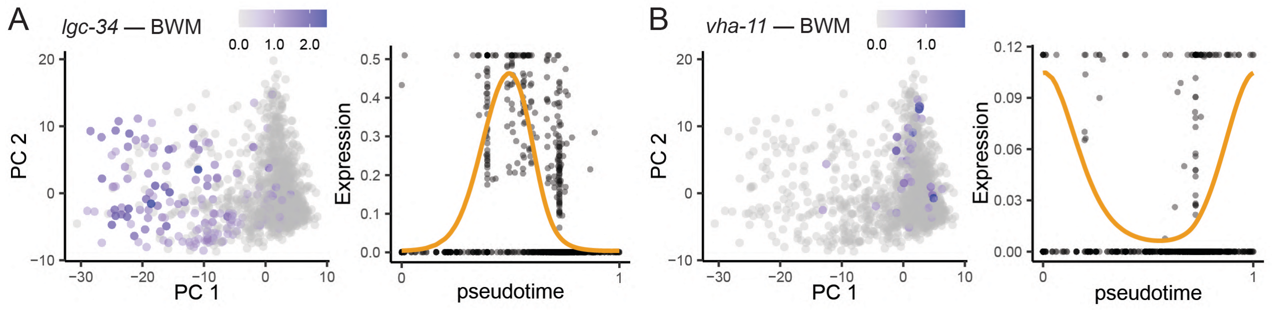


**Supporting Figure S4. Perplexity metric in a non-oscillating cell type.**

(A) Single-cell expression of *lgc-34* in body wall muscle. (B) Single-cell expression of *vha-11* in body wall muscle.

**SUPPORTING FIGURE S5**

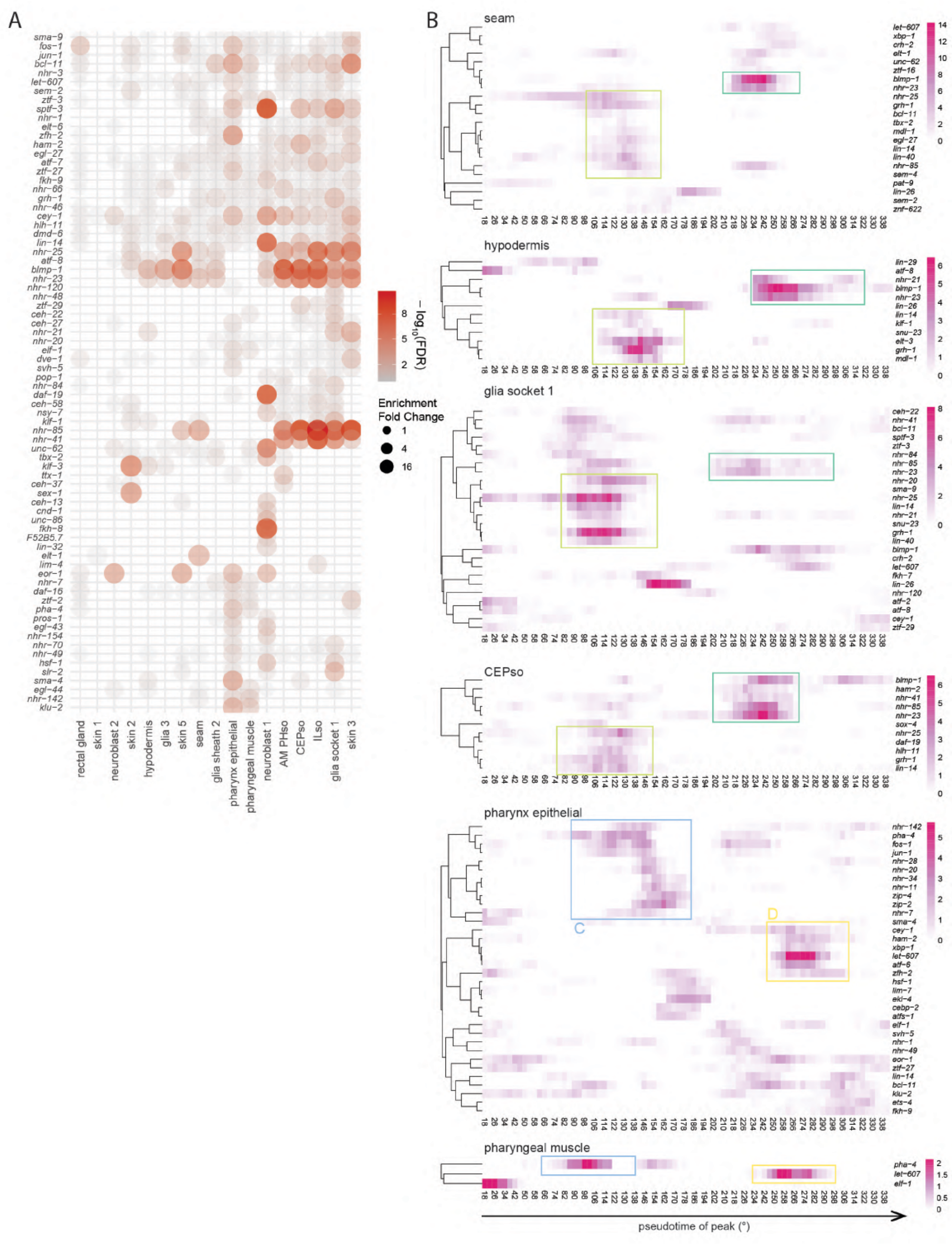


**Supporting Figure S5. Expanded analysis of TFs associated with pulsatile genes.**

(A) Dot plot representation of the Fig. 6A analysis, enabling direct identification of shared TFs. The size of the dots indicates the enrichment, the color and transparency reflect the statistical significance.

(B) TFs with enrichment in particular time bins, for selected cell types. The boxes highlight regions of interest.
